# An exploratory study of a simple approach for evaluating drug solubility in milk related vehicles

**DOI:** 10.1101/2022.10.27.514033

**Authors:** Sean Li, Justin J. Gabriel, Marilyn N. Martinez, David G. Longstaff, Martin J. Coffey, Fang Zhao

## Abstract

Milk related materials are frequently used as a vehicle for drug product administration. Therefore, drug solubility information in milk related vehicles is desirable for prediction of how they may influence *in vivo* drug release and bioavailability. However, there are very limited data published on this topic. This study explored a practical method to address the key challenges associated with solubility assessment in milk, including the sample equilibration time and cleanup procedures. Amitriptyline, acetaminophen, dexamethasone, nifedipine, piroxicam, and prednisolone were selected as model drugs to represent a wide range of physicochemical properties. Their solubilities were determined at room temperature in pH 6.8 phosphate buffer, skim milk, whole milk, reconstituted whole milk powder, and unprocessed raw milk. The overall results confirmed that milk greatly improves the solubility of poorly water-soluble drugs. However, the extent of improvement and mechanism of solubilization appeared unique for each drug, highlighting the importance of evaluating milk solubility experimentally. The method used in this exploratory study can be applied in future investigations of a broader range of drugs and milk-related vehicles.

## Introduction

Drug release from oral solid dosage forms is a critical step impacting the overall bioavailability. Milk-based drinks are often used when patients take whole capsules and tablets. They are also frequently used as liquid vehicles to disperse capsule contents or crushed tablet powder to facilitate dose administration for certain patient populations [1–3]. A variety of cow milk commodities are available to consumers, and the most common ones in grocery stores vary in the percent of milk fat from 0% (skim milk) to approximately 3.3% (whole milk) [2,4]. Components of milk are routinely used as key ingredients to manufacture a wide range of palatable nutrition products which may be used directly or after reconstitution for drug administration. Several recent studies also evaluated lyophilized milk as an excipient matrix for orally dispersible tablets which are particularly valuable for pediatric patients [5–6].

Milk contains a variety of chemical components which are expected to influence drug solubility [1–4]. Despite the frequent use of milk-based products in drug delivery, there is a paucity of primary literature data on drug solubility in these vehicles or on how that solubility effect can vary as a function of the drug’s physicochemical characteristics. Searches in PubChem and DrugBank databases revealed that most drug monographs contain only data of drug concentrations in milk from nursing mothers or dairy cows within the context of pharmacokinetic distribution and excretion to address safety concerns [7,8]. Based on the PubMed [9] search results, only one research group has published comprehensive reports on drug solubility in milk, using a flow injection serial dynamic dialysis technique [10–12]. More recently, another research group studied solubility of two model drugs in milk along with a panel of other soft foods and drinks [13–14].

There are several key challenges in performing drug solubility studies in milk due to its complex compositions and properties. Traditionally, solubility samples are agitated with excess solid drug particles at room temperature for > 4 hours (and often overnight) to reach equilibrium. However, such prolonged test procedures are not feasible for milk-related vehicles due to the potential for rapid microbial growth. Moreover, after equilibration, the excess drug particles are typically removed by centrifugation or filtration through 0.45 μm filters. This step can be problematic for milk, because it is an oil-in-water emulsion system and the aqueous phase also contains colloidal protein components. Centrifugation leads to phase separation, and 0.45 μm filters can be blocked by oil globules and colloidal protein particles. Finally, drug concentrations in saturated solution samples are typically analyzed directly by HPLC-UV methods. Milk samples, on the other hand, require proper extraction steps prior to analysis to account for all the solubilized drug in the system, including free and protein-bound drug in the aqueous phase, drug in the oil phase, and drug at the aqueous-oil interface. Additional sample treatment steps may also be necessary to reduce analytical interferences from other components in milk, such as protein.

The objective of this study was to explore a practical method to evaluate drug solubility in milk-based vehicles. Several representative vehicles and model drugs were selected and evaluated. The results provide useful preliminary data for future applications to a broader range of vehicles and drugs. The method can also be incorporated in various drug release studies to aid in drug product development activities and for the assessment of palatable, nutrient-based matrices for drug administration to special populations.

## Materials and methods

### Materials

#### Drug ingredients

Six model drugs with diverse physicochemical properties were selected for this exploratory solubility study as listed in Table 1. The source information is provided in Table 2.

**Table 1.**
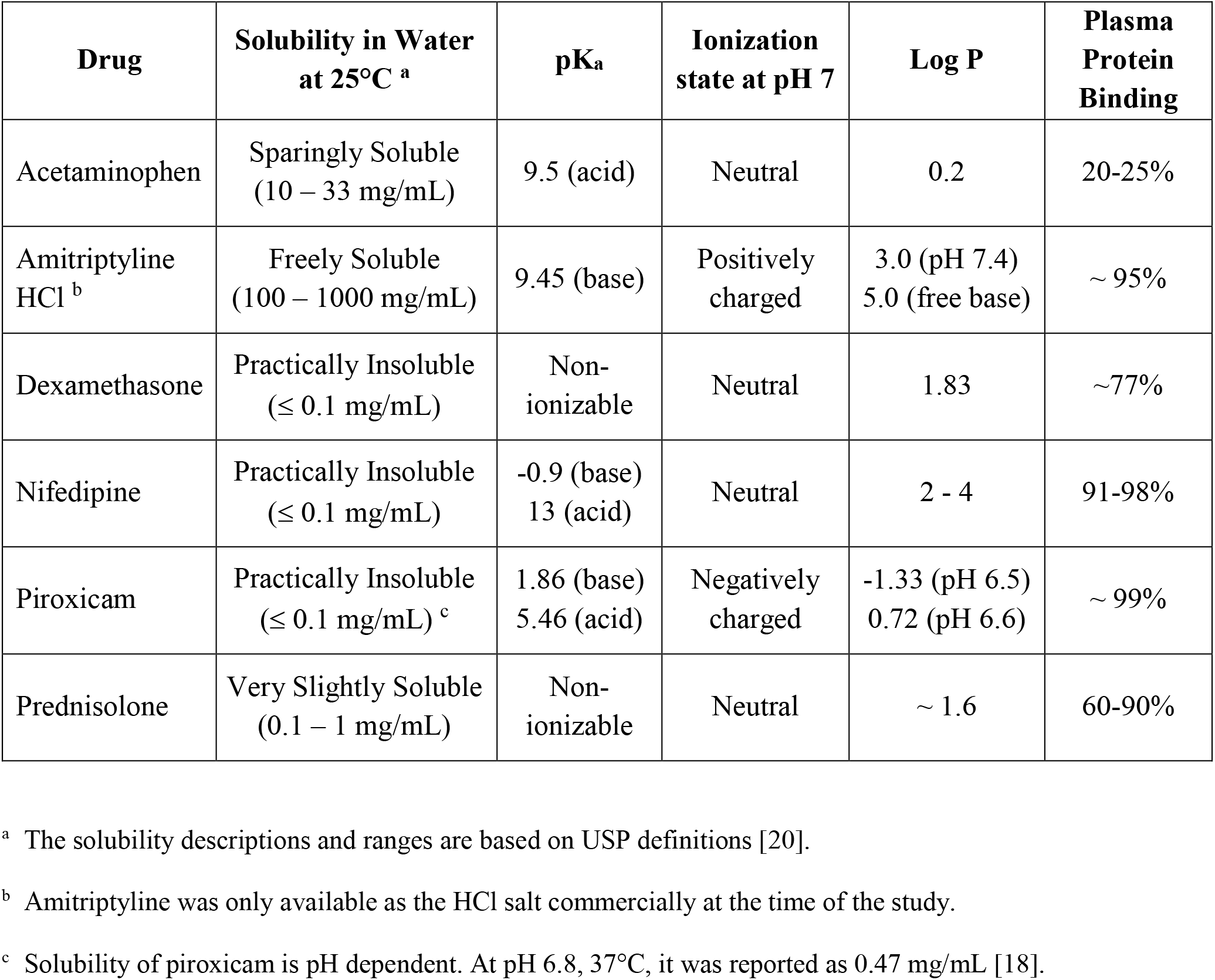
Model drugs and their selected physicochemical properties [7–8, 15–19].

| Drug | Solubility in Water<br>at 25°C <sup>a</sup> | pK<sub>a</sub> | Ionization<br>state at pH 7 | Log P | Plasma<br>Protein<br>Binding |
| --- | --- | --- | --- | --- | --- |
| Acetaminophen | Sparingly Soluble<br>(10 – 33 mg/mL) | 9.5 (acid) | Neutral | 0.2 | 20-25% |
| Amitriptyline<br>HCl <sup>b</sup> | Freely Soluble<br>(100 – 1000 mg/mL) | 9.45 (base) | Positively<br>charged | 3.0 (pH 7.4)<br>5.0 (free base) | ~ 95% |
| Dexamethasone | Practically Insoluble<br>(≤ 0.1 mg/mL) | Non-<br>ionizable | Neutral | 1.83 | ~77% |
| Nifedipine | Practically Insoluble<br>(≤ 0.1 mg/mL) | -0.9 (base)<br>13 (acid) | Neutral | 2 - 4 | 91-98% |
| Piroxicam | Practically Insoluble<br>(≤ 0.1 mg/mL) <sup>c</sup> | 1.86 (base)<br>5.46 (acid) | Negatively<br>charged | -1.33 (pH 6.5)<br>0.72 (pH 6.6) | ~ 99% |
| Prednisolone | Very Slightly Soluble<br>(0.1 – 1 mg/mL) | Non-<br>ionizable | Neutral | ~ 1.6 | 60-90% |

<sup>a</sup> The solubility descriptions and ranges are based on USP definitions [20].
<sup>b</sup> Amitriptyline was only available as the HCl salt commercially at the time of the study.
<sup>c</sup> Solubility of piroxicam is pH dependent. At pH 6.8, 37°C, it was reported as 0.47 mg/mL [18].

**Table 2.**
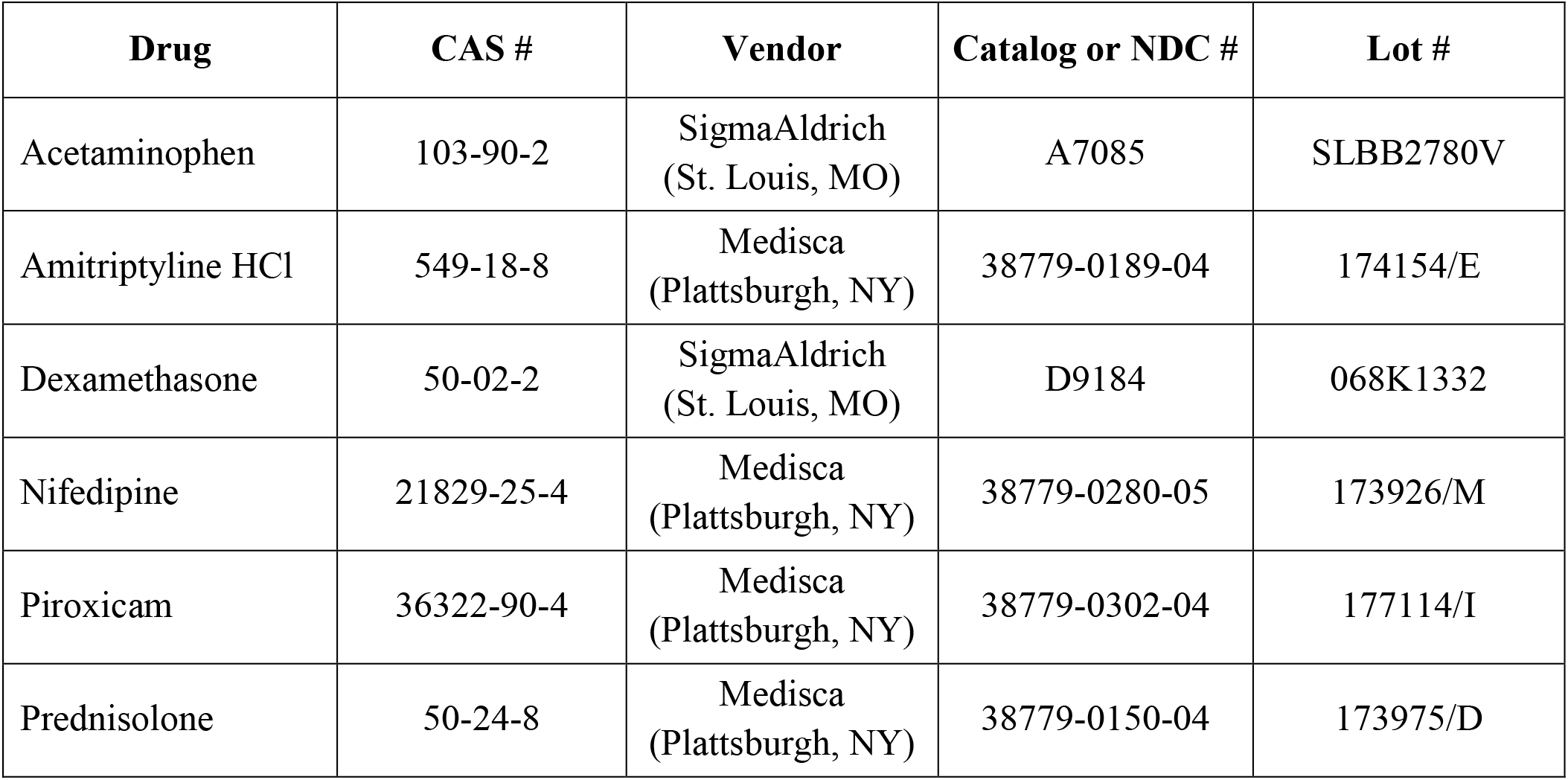
Material source information of model drugs.

| **Drug** | **CAS #** | **Vendor** | **Catalog or NDC #** | **Lot #** |
| --- | --- | --- | --- | --- |
| Acetaminophen | 103-90-2 | SigmaAldrich<br>(St. Louis, MO) | A7085 | SLBB2780V |
| Amitriptyline HCl | 549-18-8 | Medisca<br>(Plattsburgh, NY) | 38779-0189-04 | 174154/E |
| Dexamethasone | 50-02-2 | SigmaAldrich<br>(St. Louis, MO) | D9184 | 068K1332 |
| Nifedipine | 21829-25-4 | Medisca<br>(Plattsburgh, NY) | 38779-0280-05 | 173926/M |
| Piroxicam | 36322-90-4 | Medisca<br>(Plattsburgh, NY) | 38779-0302-04 | 177114/I |
| Prednisolone | 50-24-8 | Medisca<br>(Plattsburgh, NY) | 38779-0150-04 | 173975/D |

#### Milk vehicles

All milk vehicles were stored under refrigeration (2-8°C) until use. The skim and whole milk products were purchased from the local supermarket (Wegmans Food Markets, Rochester, NY) and used within the expiration dates. The fresh raw milk was provided directly by a local farm (Craft Dairy Farm, Williamson, NY) and used within 24 hours. The cows were of Holstein breed on corn and forage diet. The dry whole milk powder, Nestle Nido^®^, was purchased from Amazon or Walmart. The reconstitution was performed according to product instructions by mixing 1 oz. (31 g) powder with 8 fl. oz. (240 mL) warm (30-40°C) water. The reconstituted milk was used within 24 hours.

#### Filters

Two types of syringe filters were purchased from Phenomenex (Torrance, CA) and used for the solubility sample treatment prior to HPLC analysis. These included the Phenex-GF 1.2 μm filters (Cat# AF0-8515-12) and Phenex-RC 0.45 μm filters (Cat# AF0-3103-52).

#### Solvents for solubility sample treatment and HPLC analysis

A Milli-Q Direct 8 system from Millipore Sigma (Burlington, MA) was used to produce purified water. Acetonitrile (MeCN) and trifluoroacetic acid (TFA), both HPLC grades, were purchased from Thermo Fisher Scientific (Waltham, MA).

### Physicochemical characterization of milk vehicles

#### pH

A Mettler-Toledo (Columbus, OH) Seven Easy model pH meter was used with a gel-filled pencil-thin pH electrode from Thermo Fisher Scientific (Waltham, MA). All milk vehicles were equilibrated to room temperature prior to the pH measurement.

#### Osmolality

A μOsmette Model 5004 osmometer from Precision Systems (Natick, MA) was used for osmolality measurement, which operates based on the freezing point depression method. For each measurement, 50 μL of the vehicle was placed in the sample tube and lowered into the freezing chamber. After the solenoid-induced pulse freezing, the liberated heat of fusion was related by the microprocessor to the freezing point of the sample, and the osmolality was automatically calculated and displayed.

#### Viscosity

A DHR-2 rheometer with a Peltier temperature-controlled cup, and a double-gap concentric cylinder measurement system from TA Instrument (New Castle, DE) was used for viscosity measurements at 25°C. For each measurement, 12 g of the vehicle was placed in the cup, and the shear rate was scanned from 0.001 to 1 sec^−1^ on a log-scale with 5 points/decade. The method was set to determine the steady-state viscosity result with a maximum equilibration time of 480 sec at each point.

#### Globule size distribution

A LA-960 laser scattering particle size analyzer from Horiba (Piscataway, NJ) was used to analyze the globule size distribution of the milk vehicles at room temperature. A refractive index of 1.45 was used for the oil phase and 1.333 for the aqueous phase. The vehicle was added dropwise to deaerated water to achieve 90–95% transmission for the red laser. A circulation speed of 2 was used with no sonication, and a data acquisition of 10000 per read.

### Solubility assessment and sample treatment for HPLC analysis

Solubility assessment was performed in triplicates for each drug and milk vehicle combination. A 10 mM sodium phosphate buffer pH 6.8 was also included as a control vehicle. A general flow chart is provided in Fig 1. For each solubility sample, an excess amount of drug powder was weighed in a 20-mL glass vial. Ten mL of the vehicle, pre-equilibrated to room temperature (RT; 20-25°C), was added to the drug powder and stirred at RT for 60 minutes. It was assumed that the pH of the vehicles and the chemical stability of drugs remained unchanged during this period. Upon completion, approximately 2 mL of each solubility sample was passed through the Phenex-GF 1.2 μm filter to remove the excess solid. An aliquot of 0.5 mL filtrate was accurately transferred to a microcentrifuge tube followed by the addition of 1 mL acetonitrile (MeCN) for drug extraction and protein precipitation. The mixture was then centrifuged at 10,000 rpm for 10 minutes, and about 1 mL of the supernatant was withdrawn passed through a Phenex-RC 0.45 μm filter. The filtrate was collected in an HPLC autosampler vial and analyzed by the HPLC method described below. Placebo milk vehicles without drug were treated the same way to serve as controls for HPLC analysis.

**Fig 1.**
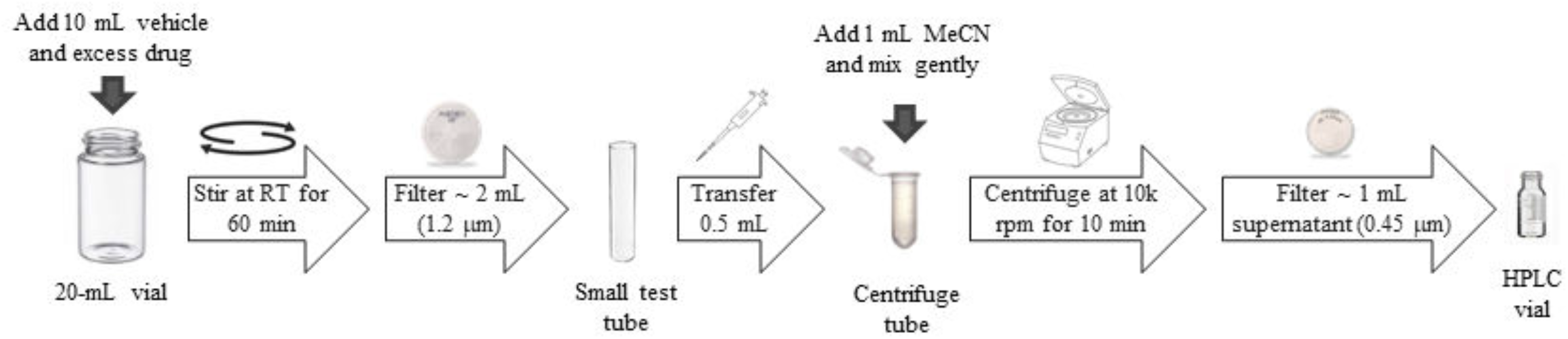
Flow chart of the solubility assessment and sample treatment steps.

Considerable back pressure was observed during the 1.2 μm filtration step of the reconstituted whole milk powder and raw milk, with or without drug. Additional investigation revealed that back pressure was inversely correlated to milk temperature. The solubility experiment was repeated at 38°C with one model drug, nifedipine, for confirmation.

### HPLC analysis

A gradient HPLC method, described in Table 3, was used to analyze the concentration of all drug solubility samples after treatment. The UV detection wavelength and sample injection volume were selected to streamline the HPLC analysis.

**Table 3.** HPLC method for drug analysis.

|  |  |
| --- | --- |
| HPLC System | Shimadzu Model# LC-2010AHT |
| Column | Phenomenex Kinetex C18, 150×4.6 mm, 5 $\mu\text{m}$, 100 Å (Phenomenex, Cat# 00F-4601-E0) |
| Column Temperature | 40°C |
| Mobile Phase | Channel A: Water with 0.1% v/v TFA<br>Channel B: MeCN with 0.1% v/v TFA<br>Linear gradient program: 5% to 95% Channel B in 10 minutes with 10 minutes of re-equilibration time.<br>Flow rate: 1.0 mL/min |
| UV Detection | 300 nm for acetaminophen<br>254 nm for dexamethasone, piroxicam, and prednisolone<br>235 nm for nifedipine |
| Inject Volume | 3 $\mu\text{L}$ for acetaminophen, dexamethasone, and prednisolone<br>5 $\mu\text{L}$ for piroxicam<br>10 $\mu\text{L}$ for nifedipine |

## Results and discussion

### Milk vehicles and properties

Milk is a biological fluid with complex chemical compositions and physicochemical properties [2, 4]. Raw and whole milk include all categories of nutrients, and the protein and lipid components are expected to have a substantial impact on the solubility of many drugs, especially the ones with poor aqueous solubilities. In addition, pH is a critical factor for solubility of drugs with ionizable functional groups.

Four milk vehicles were included in this exploratory study with the following rationales. Skim milk and whole milk are widely available at most food stores which are suitable materials for routine solubility experiments. The main difference in their lipid contents would help elucidate its effect on drug solubility. Whole milk powder provides a convenient alternative to fresh whole milk, as it has a long shelf-life and reduces the variability of whole milk due to bovine breed, environment and diet [21–22]. Raw milk is consumed in certain regions, and it is important for the development and evaluation of certain veterinary drug products intended for intramammary delivery. It is not pasteurized or homogenized, and the effect of these processes on drug solubility has not been well characterized in the past.

Several relevant physicochemical properties were evaluated for the four milk vehicles, and the results are summarized in Table 4. The pH values of all milk vehicles ranged from 6.6 to 6.8, which is consistent with published data [22]. The osmolality values of all vehicles, except reconstituted whole milk powder, were within the isotonic range of approximately 280-300 mOsm/kg. The low osmolality of the reconstituted whole milk powder was possibly attributed to the loss of some ingredients during the manufacturing process. All vehicles exhibited low viscosity at room temperature. The globule size D50 data reflected marked differences across the milk vehicles. As expected, whole milk contained much smaller oil droplets than raw milk due to the homogenization process. Interestingly, the reconstituted whole milk powder exhibited smaller D50 than whole milk. This was probably due to the presence of soy lecithin additive in the whole milk powder formulation which facilitated the emulsification process during reconstitution. Nevertheless, there were still some large globules detected in the reconstituted milk which were not reflected by the D50 result. In other words, it exhibited a wider dispersion of globule size as compared to that of whole milk. Skim milk contained very little milk fat, and the small D50 value most likely reflected the size of the colloidal protein components.

**Table 4.** Physicochemical properties of the milk vehicles.

| Vehicle | pH | Osmolality<br>(mOsm/kg) | Viscosity at 1 sec<sup>-1</sup><br>(cP) | Globule Size $D_{50}$<br>(μm) |
| --- | --- | --- | --- | --- |
| Skim milk | 6.65 ± 0.05 | 282 ± 1 | < 5 | 0.136 |
| Whole milk | 6.68 ± 0.04 | 277 ± 1 | < 5 | 0.726 |
| Reconstituted whole<br>milk powder | 6.72 ± 0.03 | 234 ± 2 | < 5 | 0.512 |
| Raw milk | 6.70 ± 0.01 | 284 ± 2 | < 5 | 3.61 |

### Model drug solubility in milk vehicles

Six model drugs were used in this study to represent a wide range of physicochemical properties as shown in Table 1. Their milk solubility experiments were performed using the exploratory method, and the data are summarized in Table 5. For all six drugs, the solubility values in the pH 6.8 buffer control were in general agreement with the published data in water. This confirms that the 60-minute equilibration time and the 1.2 μm filtration step (to remove excess drug) were suitable for regular drug powder materials.

**Table 5.** Solubility of model drugs in milk vehicles at room temperature.

| **Drug** | **Solubility (mg/mL)** |  |  |  |  |
| --- | --- | --- | --- | --- | --- |
|  | **pH 6.8 buffer**<br>**(control)** | **Skim Milk** | **Whole Milk** | **Reconstituted**<br>**Whole Milk**<br>**Powder <sup>a</sup>** | **Raw Milk <sup>a</sup>** |
| Acetaminophen | 18.6 ± 0.6 | 19.1 ± 1.0 | 18.6 ± 0.1 | 17.9 ± 0.4 | 17.7 ± 0.7 |
| Amitriptyline<br>HCl | > 20 <sup>b</sup> | > 20 <sup>b</sup> | > 20 <sup>b</sup> | Not Studied | Not Studied |
| Dexamethasone | 0.068 ± 0.001 | 0.110 ± 0.005 | 0.115 ± 0.004 | 0.123 ± 0.003 | 0.106 ± 0.001 |
| Nifedipine | 0.0051 ± 0.0001 | 0.0183 ± 0.0010 | 0.0303 ± 0.0060 | 0.0334 ± 0.0067 | 0.0120 ± 0.0014 |
| Nifedipine,<br>38°C <sup>a</sup> | Not Studied | Not Studied | Not Studied | 0.0947 ± 0.0143 | 0.0396 ± 0.0043 |
| Piroxicam | 0.287 ± 0.019 | 0.262 ± 0.005 | 0.339 ± 0.051 | 0.341 ± 0.038 | 0.389 ± 0.015 |
| Prednisolone | 0.216 ± 0.011 | 0.679 ± 0.012 | 0.667 ± 0.021 | 0.648 ± 0.011 | 0.651 ± 0.019 |

<sup>a</sup> Reconstituted whole milk powder and raw milk samples exhibited high back pressure during filtration through the 1.2 µm filters. Additional investigation revealed that the high back pressure was inherent to these two vehicles and was inversely correlated to temperature. The solubility of nifedipine was performed in these two vehicles at 38°C for confirmation.
<sup>b</sup> Amitriptyline HCl dissolved rapidly in the buffer control, skim milk, and whole milk. At > 20 mg/mL, the milk samples formed large solid clumps.

As expected, the solubilities of the four poorly water-soluble drugs were markedly improved in all milk vehicles (Table 5). However, the exact trend for each drug seemed unique and not fully rationalized by their protein binding or o/w partition characteristics (Table 1). Dexamethasone and prednisolone have closely related chemical structures with no ionizable groups. Their solubilities increased considerably in skim milk versus aqueous buffer but no additional increases in solubility occurred when tested in whole or raw milk. On the other hand, the solubility of piroxicam exhibited no difference between the control buffer and skim milk but did exhibit a substantial increase when tested in whole and raw milk. This was somewhat surprising given the high % plasma protein binding but relatively small Log P of piroxicam. For nifedipine, its solubility benefited from components in both skim and whole milk. Surprisingly, the solubility of nifedipine in raw milk was substantially lower than that in skim or whole milk. Overall the results highlighted the complexity of drug solubility behavior in milk vehicles. More experiments and data in this area are needed to enhance the overall understanding and enable accurate predictions in future.

One consistent and encouraging finding is that the whole milk powder, after reconstitution, offered similar solubility results as whole milk. Therefore, drug solubility in either vehicle can be assumed to be interchangeable and reconstituted whole milk can be considered a possible substitute in routine solubility studies. However, it was noted that the reconstituted whole milk and raw milk samples, with or without any drug, exhibited high back pressure when being filtered through the 1.2 μm filters. This was likely due to the presence of large oil globules as discussed above. When these two vehicles were warmed to > 35°C, the back pressure dropped considerably, probably due to the reduced viscosity of the oil. The solubility experiment was repeated at 38°C for nifedipine in these two vehicles without any filtration issues. The solubility increased as expected at higher temperature, and the trend was comparable for raw milk and reconstituted whole milk.

## Conclusion

Drug solubility data in milk provides valuable information to support many aspects of drug product development and administration. An exploratory method was developed to overcome key challenges associated with solubility studies in milk vehicles. The utility of the method was demonstrated in this study by six model drugs in four milk vehicles. Future studies are encouraged to apply and improve this method for the investigations of a broader range of drugs and milk-related vehicles.

## Acknowledgements

We would like to thank Mr. Eric Phillips for his assistance in the globule size analysis of the milk vehicle samples.

## Ethics Statement

This work did not involve animal use.

